# Information processing-like phenotype of human brain organoids

**DOI:** 10.64898/2026.09.09.750433

**Authors:** Hongwei Cai, Chunhui Tian, Yang Yang, Yantao Xing, Zheng Ao, Quan Wang, Hengyao Niu, Nian Wang, Omer Revah, Keith Vossel, Jungsu Kim, Orly Reiner, Mingxia Gu, Feng Guo

## Abstract

Human brain organoids recapitulate key physiological features and functions of the human brain and hold remarkable potential for studying neurological diseases. Despite clinical evidence suggesting that neurodegenerative diseases impair the information-processing ability of the human brain, the capacity of brain organoids for information processing and its relationship to neural network function remain largely unexplored. Here, we test and quantify information-processing-like property of human cortical organoids using a task-based functional phenotyping framework (Brainopheno). We demonstrate the pattern-processing-like phenotype of cortical organoids through the representation and classification of evoked neural activities in response to distinct spatial input stimulation patterns via a microelectrode array (MEA) system. Moreover, this functional phenotype emerges with network maturation and is disrupted by pharmacological perturbations, linking classification performance to the functional integrity of organoid neural networks (ONNs). Importantly, this functional phenotype also reveals functional deficits in ONNs altered by a familial Alzheimer’s disease (AD)-associated gene mutation (e.g., APP) and by monocytes from patients with sporadic AD. This work may establish a new quantifiable functional phenotype of neural organoids and provide a framework to bridge molecular and cellular profiles with neural circuit function for basic neurology, disease phenotyping, and therapeutic development.

**Significance:** Human brain organoids are widely used to study human brain circuits and neurological diseases; however, their capacity for information processing (e.g., pattern classification) remains largely unexplored. We demonstrate that human brain organoid neural networks possess experimentally quantifiable information-processing-like properties that can serve as functional phenotypes of neural network physiology and disease. Using this functional phenotyping method, we also assess functional deficits caused by a familial Alzheimer’s disease (AD)-associated gene mutation (e.g., APP) and monocytes from sporadic AD patients. This method may bridge molecular and cellular profiles with functional phenotypes of neural network physiology for basic neurology, disease phenotyping, and therapeutic development.

---

Human brain organoids, brain-like 3D *in vitro* cultures derived from human stem cells or fetal brain tissues, can recapitulate key genetic, cellular, structural, and pathophysiological features of human brain circuits^1-7^. Pioneering efforts have been made for the successful generation of cerebral organoids^8^, region-specific brain organoids^9^, and assembloids from patient-derived induced pluripotent stem cells (iPSCs)^10^, highlighting their potential for understanding and/or treating neurodevelopmental processes and neuropathological conditions^11, 12^, particularly neurodegenerative diseases such as Alzheimer’s disease (AD)^13^. For instance, brain organoids derived from AD iPSCs or lines carrying AD-risk genes (e.g., *APP, PSEN1, PSEN2*, and *APOE4*) have been generated to understand disease phenotypes and their ensuing clinical manifestations^14-20^. Moreover, brain organoids derived from healthy human stem cells have been co-cultured with microglia^21, 22^, and aged/AD monocytes^23, 24^, or exposed to various non-cellular factors, such as Aβ^25^, AD serum^26^, to model various AD-related pathologies. These organoid models have exhibited AD-associated pathologies, including Aβ and tau pathology^27^, cellular stress^28^, synaptic dysfunction^29^, neuroinflammation^30^, neurodegeneration^31^, and aberrant neural activity^32^. However, a significant gap remains between the molecular and cellular pathologies and the functional phenotype of human brain organoids, especially in the context of information processing.

The human brain performs high-efficiency information processing through functional neural circuits, which are often disrupted in neurological and psychiatric disorders^33-39^. For example, AD patients frequently exhibit deficits in information processing due to neural circuit dysfunction associated with neurodegeneration, Aβ plaque deposition, and tau pathology ^40-42^. Emerging clinical evidence further suggests that information-processing-related tasks may provide valuable opportunities for disease detection, mechanistic studies, and therapeutic development.^35, 43, 44^ Thus, it is of great interest to access and evaluate information processing phenotypes in biological neural networks within human neural organoids. However, these functional phenotypes remain largely unexplored, even for relatively basic information processing tasks. Our recent engineering advance, ‘Brainoware’^45^, established a biohybrid computing platform that leverages biological neural networks in brain organoids for AI computing, demonstrating the feasibility of basic information representation and classification using brain organoids. Among fundamental information processing functions, pattern processing represents a basic yet important task, as the human brain can perform pattern representation, classification, and more complex operations through functional neural circuits. As a proof-of-concept application, our objective is to develop an engineering strategy to access and evaluate pattern processing-like phenotype in biological neural networks within human brain organoids.

Herein, we propose the pattern processing-like phenotype of brain organoids for understanding and/or treating neurodevelopmental processes and neuropathological conditions. Specifically, we present a brain organoid phenotyping platform, or ‘*Brainopheno*’, to characterize pattern classification-based functional phenotype of cortical organoids (**Fig.1a**). Compared to current characterization of neural network functions, our Brainopheno platform may have several unique features: (1) access organoid’s basic information processing ability (e.g., pattern classification) using task-taking measurement strategies; (2) evaluate the functional phenotypes of organoid neural networks (ONNs) using quantitative metric (e.g., pattern classification accuracy); and (3) offer a new method for functional phenotyping of biological neural networks altered by different conditions (e.g., drugs, genes, cells, and other factors).

## Results

### Information processing-like functional phenotype of human cortical organoids

We proposed to characterize the pattern processing-like phenotype in human brain organoids by establishing a Brainopheno platform (**Fig.1A, Supplementary Fig.1**). After mounting onto a microelectrode array (MEA) chip, a functional human brain organoid can receive stimulation patterns and send out its response to these patterns for characterizing pattern representation, which is different from only measuring the spontaneous activity without any electrical stimulation. For instance, our platform can encode different spatial patterns (e.g., X, Y, or Z) into spatial stimulation patterns of bipolar electrical pulses. After applying different stimulation patterns to the organoid, our platform can track, record, and then process the evoked neuronal activities responding to each stimulation pattern using a simple readout function (e.g., logistic regression and principal component analysis). The representations of different stimulation patterns (e.g., “X”, “Y”, or “Z”) can be plotted into different clusters to demonstrate the pattern processing-like phenotype of brain organoids. The pattern classification accuracy can be further calculated for quantitative evaluation of this functional phenotype. Using this platform, we expect to phenotype the impairment or enhancement of organoid neural networks (ONNs) by disease genes, compounds, cells, and drug candidates, highlighting its potential for disease phenotyping, drug screening, and therapeutic development (**Fig.1B**). Based on our early work on organoid computing, we chose functional human cortical organoids for investigating pattern classification-like phenotype. These organoids were characterized with a variety of brain cell types (e.g., mature neurons, early-stage neurons, neural progenitor cells, and astrocytes), and early human brain-like structures (e.g., ventricular/subventricular zone-like structures or VZ/SVZ) to form, develop, and maintain complex 3D neuronal networks with developing neural activity and functional connectivity (**Fig.1C, Supplementary Fig.1 and 4**). Single-cell RNA-seq data further identified the cellular diversity of these organoids (e.g., intermediate progenitors, glutamatergic and GABAergic neurons) (**Fig.1D, Supplementary Fig.2**). In the next experiments using these human cortical organoids, we developed a functional phenotyping method, validated pattern processing-like phenotype, and conducted disease phenotyping.

**Fig. 1.**
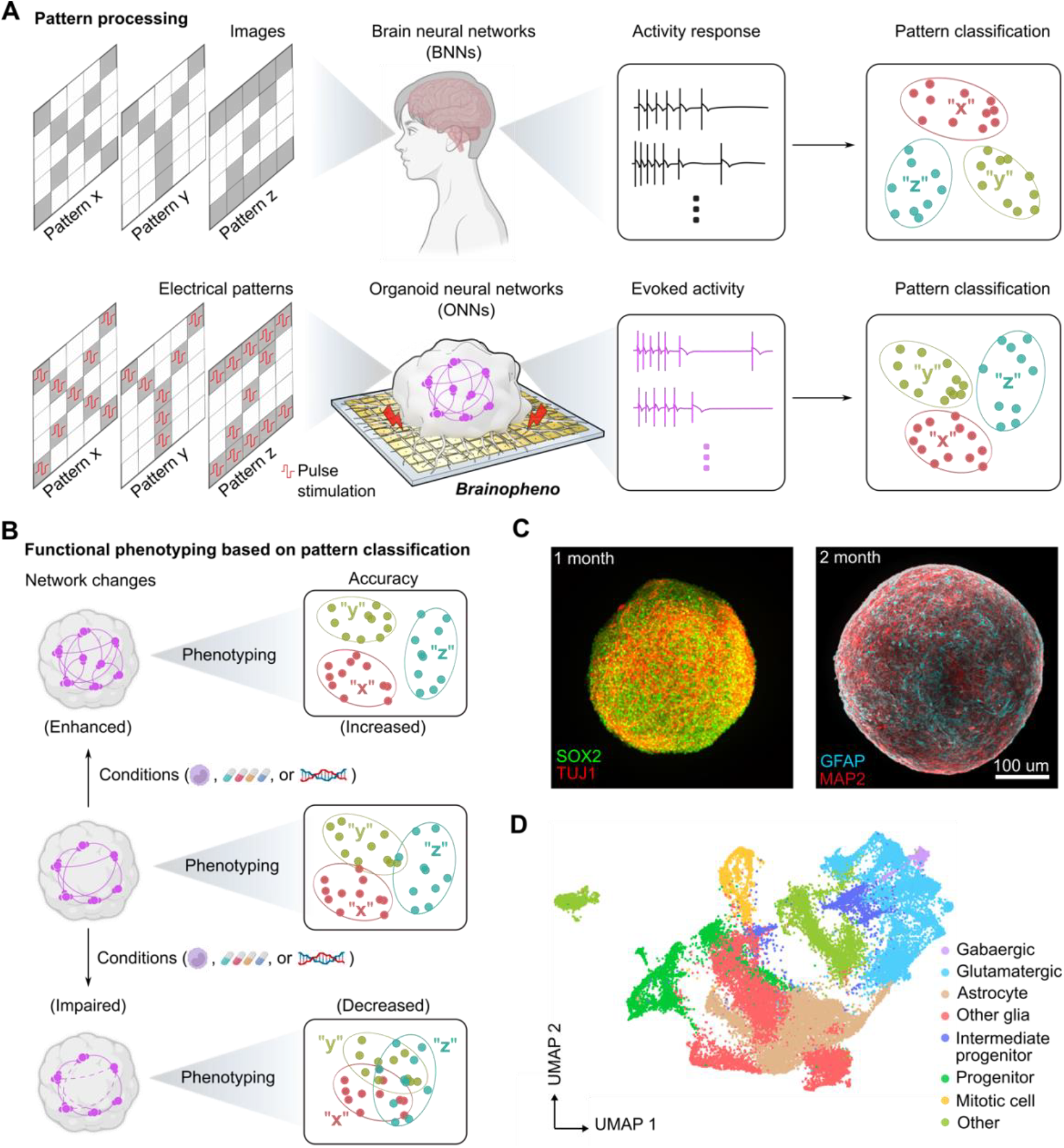
Spatial information processing-like phenotype of human brain organoids. **(A)** Schematics showing that a human brain processes pattern representation and classification, and a brain organoid exhibits the pattern processing-like phenotype using the *Brainopheno* platform. **(B)** Schematics illustrating pattern classification-based functional phenotyping of organoid neural networks impaired or enhanced by different conditions (e.g., drugs, genes, and cells). **(C)** Whole-mount immunostaining of human cortical organoids consisting of complex 3D neuronal networks with various brain cell identities (mature neuron, MAP2; astrocyte, GFAP; developing neuron, TuJ1; neural progenitors, SOX2). **(D)** Uniform manifold approximation and projection (UMAP) plots of single-cell RNA-seq results (~10,000 cells) from 2-month-old human cortical organoids showing the clusters of different brain cell types.

### The Brainopheno platform, through task-taking MEA measurements

To explore the pattern processing-like phenotype of above functional organoids, we developed a new platform and stimulation protocols of spatial patterns (**Fig.2A, Supplementary Movie 1**). We generated three different spatial stimulation patterns (e.g., *α, β*, and *γ*) and encoded the spatial information into 2D binary map of the MEA chip using bipolar voltage pulses (pulse voltage, Vpp = 500 mV; pulse duration, tp = 500 μs) and no input to present filling or no filling each pixel. To achieve pattern classification, our platform can input three different patterns *α*,*β*, and *γ* to the organoid with 30 trials of each spatial pattern, and a gap of 5 s between different trials and 30 s between different patterns. Then, our platform can record the evoked neural activity from the organoid, then analyze the organoid’s representations of different stimulation patterns (e.g., “*α*”, “*β*”, and “*γ*”) and calculate the pattern classification accuracy using a logistic regression algorithm. Applying the stimulation pattern *α* to the functional organoid mounted on the MEA chip, the representative raster plot showed the evoked electrical activities over 12 trials (**Fig.2B**, task-taking state). The distinct dynamic evoked neural activities of a 3-month-old organoid with gradual relaxation in response to the stimulation pattern *α* were different from the spontaneous electrical activities of the same organoid (**Fig.2B**, resting state without any stimulation), indicating the active display and spatial information storage. Moreover, applying different stimulation patterns to the organoid, representative raster plots (**Fig.2C**) and post-stimulation histogram (**Fig.2D**) described the distinct dynamic evoked activities of a 3-month-old organoid by different electrical stimulation patterns. Since this is a dynamic process for an organoid in responding to different patterns and trails, our platform was optimized to clearly separate these distinct spatial patterns. The pattern classification accuracy and correlated dissimilarity between different stimulation trials over time were calculated for optimizing the performance conditions (**Fig.2E**). The results showed that optimal pattern classification accuracy can be provided 30 ms after applying different stimulation patterns, which is confirmed by the maximal Euclidean distances between different stimulation pattern pairs. Using the optimized condition of our platform, the representations of each stimulation pattern were clustered into a 2D plot with each dot representing a single trial, and the distribution of each represented stimulation patterns can display the pattern processing-like phenotype of the organoid (**Fig.2F**). Correlated heatmaps of pairwise dissimilarity between different stimulation trials also confirmed the best pattern classification performance (**Fig.2F**). Next, we used the optimized Brainopheno platform to validate and characterize the pattern processing-like phenotype of functional organoids various conditions.

**Fig. 2.**
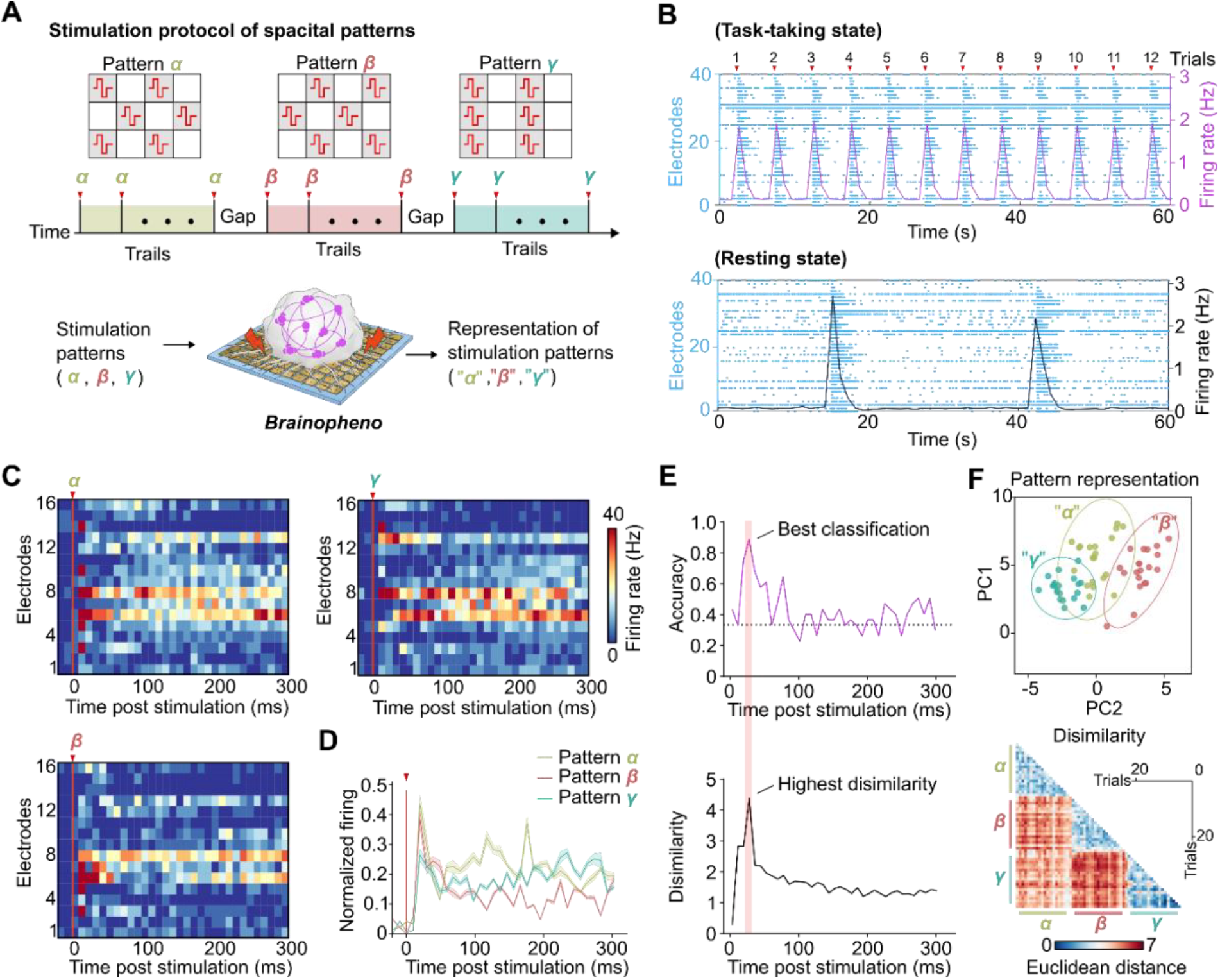
Functional phenotyping using task-taking MEA measurement. **(A)** Schematics of stimulation protocol for pattern classification-based functional phenotyping. The platform can input spatial stimulation patterns (e.g., *α, β*, and *γ*) with 30 trials of each spatial pattern (a trial interval of 5 s, and a gap of 30 s between different patterns) to the organoids, process the evoked neural activity, and analyze the representations of different stimulation patterns (e.g., “*α*,” “*β*,” and “*γ*”). **(B)** Representative raster plots of task-taking state activity (with stimulation such as 12 trials of pattern *α*) and resting-state activity (without stimulation). **(C)** Representative raster plots showing distinct dynamic evoked activities of a 3-month-old organoid by different electrical stimulation patterns. **(D)** Post-stimulation histogram of dynamic evoked activities by different electrical stimulation patterns (corresponding to **c**). **(E)** Pattern classification accuracy and correlated averaged pairwise dissimilarity over time, showing the temporal dynamics of pattern processing. **(F)** Principal component analysis (PCA) represented different stimulation patterns with each dot representing a single trial and correlated heatmaps of peak pairwise dissimilarity between stimulation trials.

### Functional phenotype association with organoid neural networks

To validate the hypothesized pattern processing-like phenotype, we tested the functional organoids using our platform. It is known that the excitation-inhibition balance in complex neural networks plays a critical role in regulating pattern separation in the human brain^46, 48^. Thus, we proposed to manipulate and validate the pattern processing-like phenotype of functional organoids by tuning global activity and excitation-inhibition balance of ONNs (**Fig.3A**). 4-month-old human cortical organoids were chosen for treatment with different neuromodulators. The ionotropic glutamate receptor antagonists, D-2-Amino-5-phosphonopentanoic acid (AP5) and 6-Cyano-7-nitroquinoxaline-2,3-dione (CNQX), were used to block/suppress excitatory transmission; the ionotropic gamma-aminobutyric acid (GABA_A_) receptor competitive antagonist bicuculline was used to block inhibitory transmission; the sodium channel inhibitor, tetrodotoxin (TTX), was used to block firing in excitatory and inhibitory neurons. We compared the resting-state and task-taking MEA measurements of these organoids before and after treating them with the above modulators for 20 mins, respectively (**Fig.3B, Supplementary Fig.3**). We found that the pattern classification accuracy was significantly decreased from 89% to 55.5% after treating with AP5 + CNQX, while the same organoids were characterized with reduced neural activities and impaired functional connectivity after the same AP5 + CNQX treatment. Moreover, our results also showed that the pattern classification accuracy was significantly decreased from 88.5% to 78% after treating with bicuculline. The same organoids were characterized with increased neural activities and similar functional connectivity after the same bicuculline treatment, which is consistent with other reports^1^. Furthermore, after treating with TTX, we found these organoids had almost lost their pattern classification accuracy from 85.5% to 32.5% (random baseline). Meanwhile, the same TTX-treated organoids lost spontaneous activity and functional connectivity, indicating the complete loss of their network function.

**Fig. 3.**
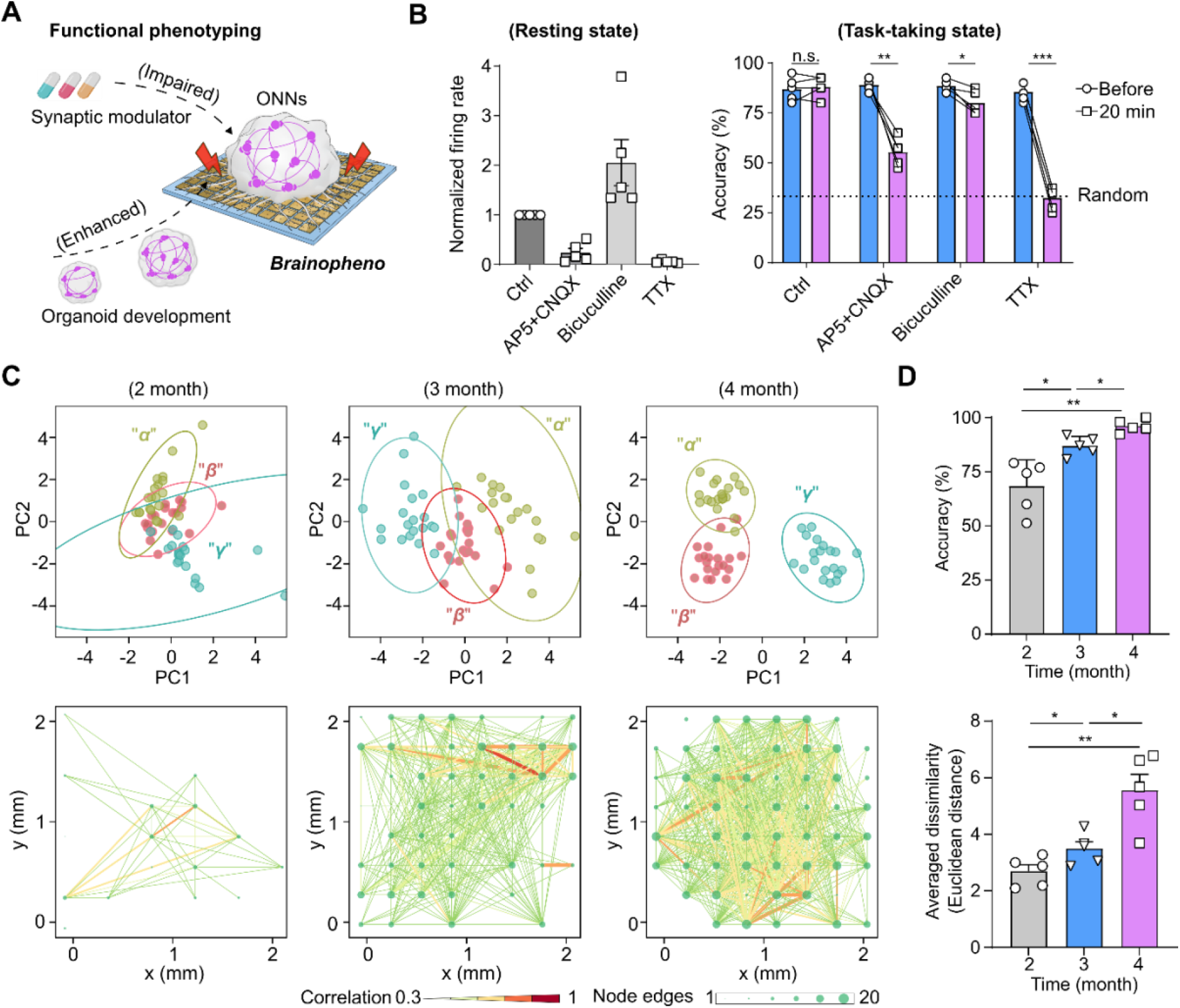
Functional phenotype association with organoid neural networks. **(A)** Schematics illustrating the pattern processing-like phenotypes of impaired or enhanced organoid neural networks (ONNs) through synaptic modulator regulation or organoid development, respectively. **(B)** Normalized mean firing rate changes (resting state) and pattern classification accuracy changes (task-taking state) of 4-month-old human cortical organoids before (Ctrl) and after a 20 min treatment with synaptic modulators including AP5 + CNQX (glutamate receptor antagonists), Bicuculline (GABAA antagonist), and TTX (sodium channel blocker), respectively (mean ± s.e.m., n = 5 organoids, 3 independent experiments). The dashed line indicates the random baseline for 3-class classification. **(C)** Representative pattern classification-based functional phenotypeand functional connectivity maps of the developing ONNs from months 2 to 4. **(E)** Quantification of pattern classification accuracy and averaged dissimilarity of the developing ONNs (mean ± s.e.m., n = 5 organoids, 3 independent experiments).

Along with the excitation-inhibition balance of ONNs, we also considered the potential enhancement of pattern processing-like phenotype upon the development of ONNs, since it is known that the brain organoids improve their network function during organoid development^1, 56^. Thus, we proposed to characterize the association of pattern classification with organoid development (**Fig.3A**). During the 4-month organoid development and maturation, we found the maturation and improvement of functional ONNs in these cortical organoids (**Supplementary Fig.4**). Using our platform, the pattern classification of these developing organoids was characterized and calculated. Our representative results showed the three merged patterns started to cluster and distinguish from each other from month 2, to 3, and to 4, and functional connectivity maps of the developing organoids also indicated the significant enhancement of the functional network (**Fig.3C**). Moreover, our quantification results also showed the significant increase of pattern classification accuracy and averaged dissimilarity of these developing organoids from month 2 to 4 (**Fig.3D, Supplementary Fig.5**). Based on these associations of pattern classification with functional ONNs, we proposed functional phenotyping of ONNs influenced by factors such as molecules, genes, and cells.

### Functional phenotyping of the familial Alzheimer’s gene mutant

Understanding the intricate relationship between genotype and phenotype is of great importance for basic biology and translational medicine. We thought our platform might facilitate the functional phenotype of disease-associated gene mutation. For example, it is known that the amyloid-β precursor protein (*APP*) gene mutation causes familial Alzheimer’s disease (AD), characterized by amyloid-β deposition and cognitive decline^57, 58^. So far, brain organoids from iPSCs derived from familial patients carrying the APP gene mutation have been developed for modeling cellular and molecular features of AD pathology^31, 59-61^. However, their phenotype in recapitulating or mimicking cognitive decline and function loss in AD patients is still largely understudied. Thus, we proposed functional phenotyping of familial Alzheimer’s gene (*APP* mutation) in cortical organoids using our platform, highlighting its impact on AD pathology and translational medicine. Following the same protocol, human cortical organoids were generated from iPSCs derived from a familial Alzheimer’s disease patient carrying a duplication of the *APP* gene^62^ and a healthy control (HC) donor. The 3-month-old *APP* and HC organoids were characterized using immunostaining with Aβ markers (4G8 and 6E10). Representative Aβ immunostaining (**Fig.4A**) and quantification of Aβ plaques (**Fig.4B**) show robust Aβ deposition in *APP* organoids, whereas HC organoids display little to no deposition. The 3-month-old *APP* and HC organoids were further characterized using the resting-state and task-taking MEA measurements. Our results showed the APP organoids had similar mean firing rate and global efficiency compared to the HC organoids (**Fig.4C**), while they showed significantly different pattern classification results. First, the 3-month-old *APP* organoid showed different evoked activity than HC organoids during task-taking MEA measurement with a trial of stimulation pattern *α* (**Fig.4D, Supplementary Fig.6C**), indicating reduced classification of evoked representations. Moreover, PCA plots showed the APP organoid displayed a merged pattern compared to the HC organoids, which is consistent with their functional connectivity maps (**Fig.4F**). Then, quantified results also confirmed that the *APP* organoids had lower pattern classification accuracy and averaged dissimilarity compared to the HC organoids. Furthermore, to model the impact of soluble toxic amyloid-β on healthy brain tissues, we also demonstrated the affected pattern classification of HC organoids by soluble oligomeric amyloid-β-42 (**Supplementary Fig.6A, B**). Thus, using our platform, we determined the impaired pattern classification by a familial Alzheimer’s gene mutation (*APP*) and its soluble toxic amyloid-β product.

**Fig. 4.**
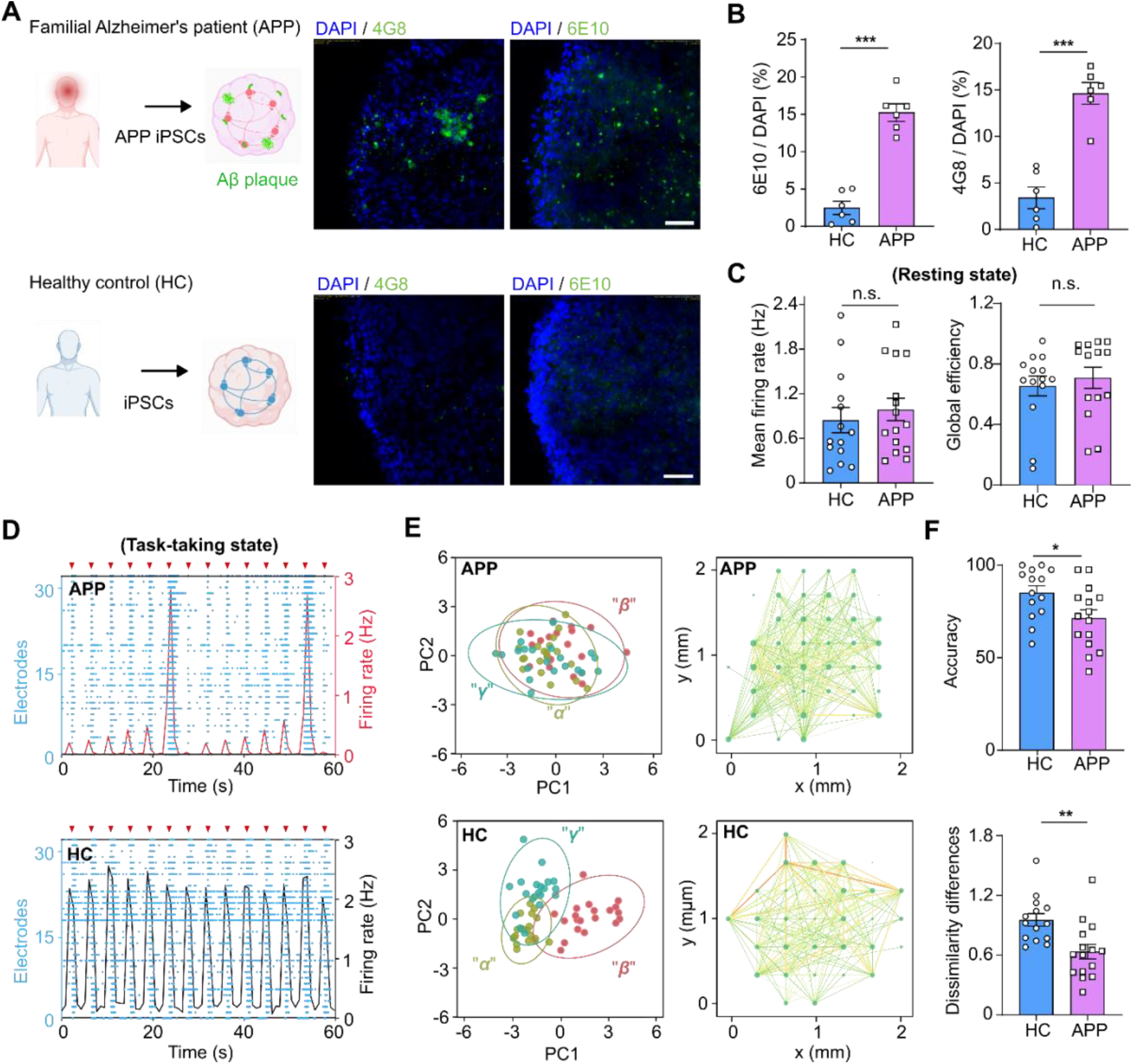
Functional phenotyping of gene mutation. **(A)** Generation of human cortical organoids from iPSCs derived from a familial Alzheimer’s disease patient carrying the amyloid precursor protein (APP) gene mutation and a healthy control (HC) donor. Representative staining of Aβ markers (4G8 and 6E10) of 3-month-old APP and HC organoids, highlighting amyloid-beta deposition in the APP organoids. **(B)** Quantification of Aβ plaques in these organoids (mean ± s.e.m., n = 6 organoids, 3 independent experiments). **(C)** Quantitative resting-state activities of 3-month-old APP and HC organoids, including mean firing rate and global efficiency (mean ± s.e.m., n = 15 organoids, 3 independent experiments). **(D)** Raster plots showing the representative evoked activity of 3-month-old APP and HC organoids during task-taking MEA measurement. **(E)** Representative pattern classification-based functional phenotype (left) and functional connectivity map (right) of 3-month-old APP and HC organoids. **(F)** Quantified pattern classification accuracy and averaged dissimilarity of 3-month-old APP and HC organoids (mean ± s.e.m., n = 15, 3 independent experiments).

### Functional phenotyping of neuroimmune interaction

Next, we validated the feasibility of our platform for functional phenotyping of neuroimmune interaction in brain organoids. It has been intensively reported that the immune system shapes and supports the function of brain neural networks in humans and plays an important role in brain aging and disease^63-65^. Moreover, it is known that circulating monocytes may show protective and healing properties or promote disease progression after infiltrating into the brain^66-68^. To understand the interaction of circulating monocytes with healthy brain tissues in aging and Alzheimer’s conditions, we demonstrated the pattern processing-like phenotyping of healthy organoids influenced by circulating monocytes from various donors and patients. The healthy 2-month-old human cortical organoids were cocultured with the monocytes from AD patients (above 65 years), old (age-matched controls of AD), and young (20~29 years) donors, respectively. After coculturing for 2 days, these monocyte-infiltrated organoids were plated onto the MEA plates. Followed by 1 month of culture on MEA, these organoids were ready for the resting-state and task-taking MEA measurements, respectively (**Fig.5A, Supplementary Figs. 7, 8**). The PCA plots (**Fig.5B**) showed the representative pattern classification of these organoids without any treatment (Ctrl) and cocultured with the monocytes from AD patients (AD), old age-matched controls of AD (Old), and young (Young) donors, and their pattern classification accuracies were calculated (**Fig.5C**), respectively. Compared to the Ctrl group, we found that (1) the AD group showed an obviously impaired functional phenotype; (2) the old group had a slightly improved functional phenotype; and (3) the Young group had asignificantly improved functional phenotype. Meanwhile, the representative functional connectivity maps of these organoids at different coculture groups also reflected the corresponding network functions (**Fig.5B**). It seems that the AD monocytes may induce neuroinflammation and neurodegeneration in the neural networks of healthy cortical organoids. Moreover, the young and old monocytes both can support the functional neural networks of healthy cortical organoids, where the old monocytes may become pro-inflammatory and lose some of their neural supportive function compared to the young monocytes. Single-cell RNA sequencing was performed on organoids from different coculture groups to assess how various monocyte populations affect the neural networks of healthy cortical organoid (**Fig.5D, Supplementary Fig.9**). The bar plots show the gene ontology (GO) terms associated with upregulated and downregulated neuronal genes in the ‘AD vs Ctrl’, ‘Old vs Ctrl’, and ‘Young vs Ctrl’ comparisons, which are consistent with our functional phenotyping results. These findings demonstrate that ONN functional phenotyping can capture both immune cell–mediated improvements and impairments, underscoring its potential for studying neuroimmune interactions and for testing novel therapeutic approaches in brain disease models.

**Fig. 5.**
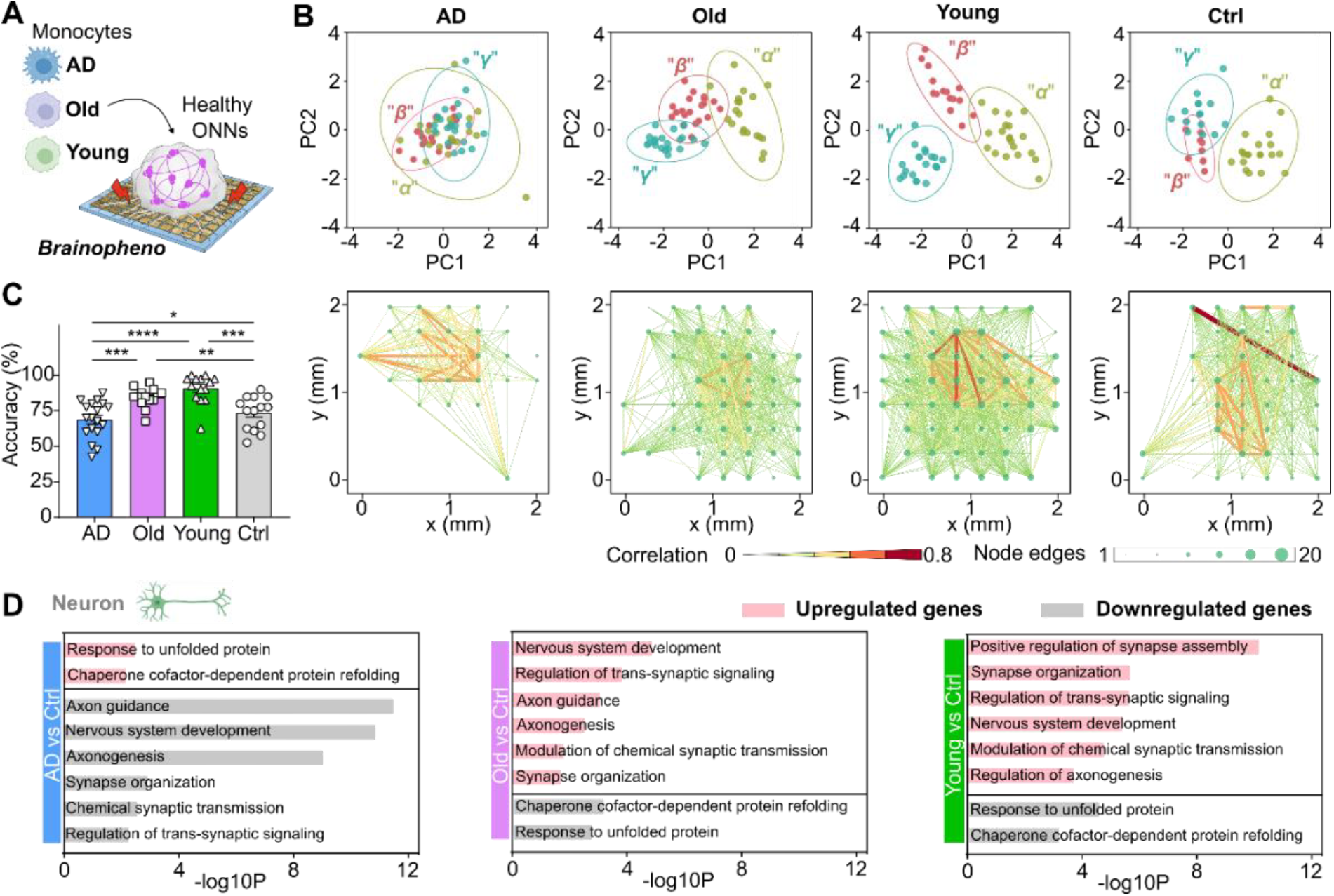
Functional phenotyping of neuroimmune interaction. **(A)** Schematic illustrating the use of *Brainopheno* for functional phenotyping of healthy 3-month-old human cortical organoids cocultured with the monocytes from AD patients (above 65 years), old (age-matched controls of AD), and young (20~29 years) donors, blank controls without monocytes (Ctrl). **(B)** Representative pattern classification-based functional phenotypes and functional connectivity maps of 3-month-old healthy human cortical organoids treated with AD monocytes (AD), old monocytes (Old), young monocytes (Young), and without monocytes (Ctrl). **(C)** Quantification of pattern classification accuracy at different conditions (mean ± s.e.m., n = 14, from 3 independent experiments). **(D)** Bar plots showing the gene ontology (GO) terms of upregulated genes and downregulated genes of neurons in the ‘AD vs Ctrl’, ‘Old vs Ctrl’, and ‘Young vs Ctrl’ groups.

## Discussion

We have developed a Brainopheno platform that enables functional assessment of spatial information processing in cortical organoids. As a proof-of-concept demonstration, our platform can access a pattern processing-like phenotype of cortical organoids using task-taking MEA measurements. This spatial information processing-like phenotype is validated by correlating quantitative accuracy with the functionality and integrity of ONNs, highlighting the innovative circuit-level computational capacity of brain organoids. We further applied the platform to evaluate the impairment of ONNs by a familial AD-associated APP mutation, by soluble Aβ molecules, and by interactions with monocytes from sporadic AD patients. Our findings not only show that brain organoids can capture molecular and cellular hallmarks of disease, but also offer new functional phenotypes that may better reflect the clinical dysfunction of AD patients. Thus, our platform may provide a unique solution to bridge cellular and molecular pathology with circuit-level functional phenotype, shining light to interrogate human neural network physiology, establish disease-relevant computational phenotypes, and accelerate therapeutic development.

There are several opportunities to further expand both the capabilities and applications of the Brainopheno platform. Importantly, the task-based strategy is not limited to electrical stimulation and MEA recordings. The system innovation can be achieved by replacing current information input using optogenetic or ultrasonic stimulation, and by advancing readouts using calcium imaging and high-density probes (e.g., Neuropixels), enabling deeper interrogation of ONN dynamics across spatial and temporal scales. The current Brainopheno platform has only demonstrated spatial information processing, such as pattern processing. Beyond spatial information processing, the platform can be extended for temporal and temporospatial information processing tasks, showing the potential to bring organoid phenotyping closer to capturing the computational repertoire of in vivo cortical circuits. Furthermore, along with cortical organoids, other brain region–specific organoids (e.g., hippocampal organoids, thalamic organoids) and/or assembloids derived from patient and healthy iPSCs can be applied for characterizing their information processing-like phenotype, highlighting the translational impact ranging from memory impairment to sensorimotor dysfunction across various disease conditions.

Despite these advances, some challenges remain for the widespread application of the Brainopheno platform. Current MEA stimulation and recording only interact with the organoid surface in contact with rigid electrodes, limiting access to the full 3D neural networks of the organoid^69^. Emerging approaches, including embedded 3D or flexible electrodes, and multiphoton imaging, may provide more comprehensive interrogation of network dynamics^7, 12^. Organoid heterogeneity also remains a significant limitation. Although the variability can be partially mitigated by selecting organoids with robust early-stage activity, engineered solutions such as vascularized or perfused organoids may yield more consistent and physiologically mature networks, thereby improving reproducibility and robustness. In addition, the limited diffusion of drugs and immune cells into spherical organoids constrains their use in modeling systemic interactions. Innovations in perfusion microfluidics, vascular engineering, and adaptive co-culture systems may help overcome these barriers^70, 71^, ultimately enhancing the translational potential of organoid-based functional phenotyping. Although we have demonstrated basic pattern classification-based functional phenotyping of brain organoids, it remains challenging to assess more advanced information-processing–like phenotype in brain organoids, as which may require higher-order network properties and circuit organization^51, 52^ that current organoids may not fully recapitulate. In addition, important ethical considerations and technical challenges must be carefully addressed when attempting to access advanced information-processing phenotypes in human brain organoids.

## Materials and methods

### Human cell resources

The human ESCs (WA09), AD APP duplication IPSCs (UCSD241i-APP2-3) were obtained from the WiCell Institute. The healthy donor IPSCs (051179) were obtained from the New York Stem Cell Foundation. We handled these cells following the guidelines from the WiCell Institute, the New York Stem Cell Foundation, and the Biosafety Committee of Indiana University. The stem cells were maintained in mTESR plus medium (Stemcell Technologies) on GFR Matrigel (Corning) coated 6-well plates in a humidified chamber at 37ºC and 5% CO2. Medium change was performed every other day. The cells were passaged every 6-8 days using ReLeSR™ passaging reagent (Stemcell Technologies).

### Stem cell culture and cortical organoid generation

Cortical organoids were generated following the other protocol that we adapted in the lab^45^. Briefly, the stem cell line was fed every other day with mTeSR+ medium and passaged every 6 days at 70%-80% confluence. The stem cell colonies were dissociated using ReleSR for 6 minutes at room temperature and centrifuged for 3 minutes at 200 g. The embryonic bodies (EBs) were generated by plating 9,000 cells per well in 96-well u-bottom spheroid plates and centrifuging for 2 minutes at 100 g. The cells were kept in 96-well plates for 30 days, with media changes every other day. Then, the organoids were transferred to 24-well ultra-low binding plates and kept in suspension under orbital shaking (60 rpm). The cells were kept in a humidified chamber at 37 °C, supplied with 5% CO_2_. Detailed medium composition could be found in **Table S1**.

### Immunofluorescence staining

Cortical organoids were fixed in 4% PFA (paraformaldehyde) in PBS at 4 °C overnight, followed by 30% sucrose incubation for 1 day. Then the organoids were embedded into Tissue-Tek O.C.T. Compound and sectioned to 25μm slices in a cryostat. The slices were first processed with heat-induced antigen retrieval (20 minutes in citrate buffer at 95 ° C) and followed by a 2-hour blocking solution (0.3% Triton X-100 and 5% normal donkey or goat serum in PBS) incubation at room temperature, then incubated with primary antibodies in block solution overnight at 4 °C. Next, the samples were washed three times with PBS and incubated with corresponding secondary antibodies in blocking solution for 2 hours at room temperature. After more than 3 times of washing, the samples were stained with DAPI solution (1 μg/mL) and mounted using ProLong Gold antifade reagent. Detailed antibody information can be found in Table S2.

### Whole-mount staining of organoids

The organoids were washed with PBS 3 times and fixed in PFA for 1 hour, and the fixed organoids were washed with PBS for another 3 times. Then, the samples were incubated in blocking solution (1x PBS containing 0.5 % Tween 20, 0.5 % Triton-X100, 1 % BSA, 3 % FBS, and 0.01 % (wt/vol) sodium deoxycholate solution) for 2 hours at room temperature and then transferred to primary antibody diluted in blocking solution at 4 ºC for 2 days. Followed by 5 times wash with 1x PBS and secondary antibody incubation at 4 ºC for another 2 days. The samples were placed on a gentle rocker during antibody incubation. And the labeled samples were conducted a serial dehydration using serial Ethanol solutions (50%, 70%, 100%, 100%) for 2 hours each, and the dehydrated organoids were transferred to Benzyl Alcohol/Benzyl Benzoate (BABB) clearing solution (benzyl benzoate: benzyl alcohol = 2:1). And the cleared organoids were imaged by a confocal microscope (Olympus OSR spinning disk) using 20x objective.

### Sample preparation for single-cell sequencing

Single-cell samples were dissociated using Papain-based Dissociation (Miltenyi Biotec) following the manufacturer’s instructions with slight modification. In brief, organoids were first cut into two hemispheres and washed three times in HBSS (without Ca^2+^ and Mg^2+^, −/−) to wash out debris and culture medium. The organoid pieces were then incubated in Enzyme P (enzyme mix 1) in a 6-well plate for 15 minutes, followed by adding Enzyme A (enzyme mix 2) for another 30 minutes. During Papain incubation, the mix was placed on an orbital shaker at 90RPM and triturated using a wide-orifice 1,000-ml tip every 10 minutes. The cells were then filtered through a 30-μm filter and washed with organoid medium at 4 °; C, centrifuged for 10 minutes at 300 g and 4 °C. The cells were washed with organoid medium at 4 °;Cfor 3 times, analyzed and counted using Trypan Blue assay, and resuspended to a desired concentration (>1 million/ml)

### Single-cell RNA-seq data processing

Raw FASTQ file of individual samples generated from 10x Chromium scRNA-seq were preprocessed and aligned using 10x cloud (10x Genomics Cell Ranger (v6.1.2), referred to human reference hg38 (10x Genomics refdata-gex-GRCh38-2020-A)). 10x Cell Ranger filtered data were used for downstream QC assessment, and filters were then applied to keep cells with at least 500 genes, less than 100,000 unique molecular identifiers (UMIs), and less than 25% UMIs mapped to mitochondrial genes. Downstream analyses were performed in R v4.1.0 and Python 3.

### Plating organoids onto MEA plates

Cortical organoids at different ages (from one month to four months) were plated in 6-well MEA plates with one organoid per well. Each MEA plate well has 64 low-impedance platinum electrodes (30 μm in diameter with 200 μm spacing, 8×8 array), yielding 384 channels in total. The MEA plates were first coated with 0.05% polyethylenimine (PEI) solution for 1 hour and 10 μg/mL laminin for another 1 hour, and then the organoids were plated into them. The organoids were fed every other day with the medium in **Table 1**. The spontaneous activity of the organoid was measured 2-6 hours after medium change. The extracellular recording was conducted using a Maestro Edge MEA system and Spontaneous Neural configuration in the Axis Navigator software following the manufacturer’s guidelines (Axion Biosystems). The organoids were placed in the MEA system for 2 minutes to allow the environment to settle down (37 °;C and 5% CO_2_) and then recorded for 5 minutes.

### Spontaneous activity analysis

The neural activity spikes were detected by Axis Navigator software using an adaptive threshold algorithm (threshold to 5.5x of the standard deviation of the background noise for each channel). The gain was set to 512x and filtered with a 300 Hz high-pass filter. The raster plot of the detected spikes was created using the Neural Metrics Tool provided by Axion Biosystems. The quantification of data (burst frequency, mean firing rate, network bursts) was conducted by using the internal processing module of Axis Navigator software.

### Functional connectivity via spike time tiling coefficient

Correlation of neural activity spike trains from each channel was evaluated using the spike time tiling coefficient (STTC)^72^. A publicly available Python function (Elephant^73^) was used to compute the STTC. The correlation time window (Δt) was set to 20 ms to compute the functional connectivity weights of all channels. And the correlation matrix was further thresholded at 0.3 for further analysis (such a threshold could rule out most of the random connectivity). A customized Python script was developed to draw the functional connectivity map based on the calculated correlation weights of all channels. The nodes and connections were drawn corresponding to the physical location of the electrodes. And the network indices (e.g., efficiency, clustering, small-worldness) were analyzed using a publicly available Python library (Networkx^74^).

### Electrical stimulation and decoding

The plated cortical organoids were stimulated with 2D-patterned voltage bipolar pulses using the MEA system. The patterned stimulation was given as shown in the main figures with pulse voltage = 500 mV and pulse time = 500 μs. The four different patterned stimulations were repeated 30 times each with 5 5-second intervals, and 30 30-second intervals when switching patterns. The evoked electrical activity of the organoids was recorded and subjected to further analysis. The evoked activity from the top-20 most active channels in the spontaneous recordings was selected, and neural activity spikes were detected using the same 5.5x adaptive threshold as in the spontaneous activity processing. The spikes were binned in 10-ms time bins to a 2D matrix (x=channel, y=time, value=total spikes of one channel in the 10-ms time bin), and the spikes from the first 10 ms after stimulation were removed to avoid artefacts induced by stimulation. The 2D matrix was then split into a 2:1 (train: test) ratio to train a logistic regression algorithm to decode the neural activity, and the pattern recognition accuracy was obtained from test data decoded by the trained logistic regression algorithm.

### Principal component analysis (PCA) of evoked neural activity

We developed a custom Python script to perform PCA (Scikit-learn^75^) to visualize the evoked neural activity patterns by different spatial electrical stimulation. The 2D matrix representing evoked neural activity was first windowed at [-200 ms: 200 ms], and was then linearly transformed to a 1D array. The 1D array was then input to the PCA algorithm to reduce the dimension to 7 principal components, and the neural activity patterns were visualized by plotting each stimulation-evoked activity trail in the space of the first two components. While the neural trajectories video was generated by performing PCA for neural activity in each time bin, we could get a data point of each stimulation-evoked activity trail at every 10 ms time bin and get a video to visualize the neural trajectories representing different spatial stimulation over time.

### Drug treatment and MEA measurement

The neural modulation drug experiment was performed with organoids adhered to 6-well MEA plates (3-month organoids, n=5 for each individual drug) using the drugs as follows: 25 μM CNQX+25 μM AP5, 20 μM bicuculline, 50 μM muscimol, 30 μM baclofen, and 1 μM TTX. In this experiment, baseline results (including spontaneous and evoked activity) were obtained 2-6 hours after medium change, and the treated results were obtained 20 minutes after the compound administration (half medium change with 2x desired concentration). After the experiment, the drug was removed by three times of PBS washing and fresh medium replacement.

### Monocyte isolation from human PBMCs

The human PBMCs were purchased from Stemcell Technology, where they obtain the cells using Institutional Review Board (IRB)-approved consent forms and protocols. Monocytes were isolated from PBMCs from six different healthy old (n=3, aged 60 +) and young (n=3, age < 30) donors. We employed the Classical Monocyte Isolation kit (Miltenyi Biotec) to isolate monocytes according to the manufacturer’s guidelines using the LS Column and the corresponding magnet (Miltenyi Biotec). Typically, around 1.5 × 10^^5^ monocytes could be isolated from 1 million PBMCs.

### Monocytes coculture with cortical organoids

Two-month cortical organoids (in suspension) were first tested by MEA to validate their functions (active electrode > 6, mean firing rate > 0.2 Hz) and then cocultured with 1 × 10^^5^ monocytes per organoid in an ultra-low attachment 96-well plate in 300 μl cortical organoid medium. After two days, the organoids were transferred to MEA plates for another 1-month culture to let the organoids fully attach to the MEA and subjected to spontaneous and evoked electrical activity measurement. The signal was analyzed using the method mentioned above.

### Statistical information

The statistical analysis between groups was conducted using GraphPad Prism 7. T-test (two groups) and ANOVA (more than two groups) were used to calculate the significance between groups, denoted as ^*^: p < 0.05, ^**^: p < 0.01, ^***^: p < 0.005,^****^: p < 0.001.

## Data availability

Single-cell RNA-seq data have been deposited at GEO (Accession number: GSE310391, reviewer token: kzkbgqeavdcbziz) and will be publicly available as of the publication date. Any additional information required to reanalyze the data reported in this paper is available from the lead contact upon request.

## Acknowledgments

F.G. wants to acknowledge the support from the National Science Foundation Award (EFRI2422149).

## Author contributions

F.G. and H.C. conceived the study and designed experiments. H.C., C.T., Z.A., and Q. W. performed the experiment. H.C., Y.Y., C.T., Y. X., H. N., N. W., O. R., J. K., and M.G. analyzed the data. F.G. and H.C. wrote the manuscript. All authors read and provided feedback on the manuscript.

## Notes

### Competing Interest Statement

The authors have declared no competing interest.

