## Supplementary material for "Information processing-like phenotype of human brain organoids": SI

**Supporting information:**

**List of Contents**

**Supplementary Figures**

Supplementary Fig. 1 Workflow of functional phenotyping of brain organoids based on pattern classification.
Supplementary Fig. 2 Single cell RNA seq data
Supplementary Fig. 3 Characterization of organoid function changes by molecular neuromodulators.
Supplementary Fig. 4 Maturation of functional neural networks in human cortical organoids.
Supplementary Fig. 5 Functional phenotyping of developing cortical organoids.
Supplementary Fig. 6 Functional phenotyping ONNs impacted by Aβ
Supplementary Fig. 7 Monocytes impact organoid functional networks
Supplementary Fig. 8 Organoid phenotype impacted by young, old, and AD monocytes.
Supplementary Fig. 9 Additional single-cell seq data of monocyte-cocultured cortical organoids

**Supplementary Movies**

Supplementary Movie 1. The pattern classification process on the Brainopheno platform.

**Supplementary Tables**

Supplementary Table 1. Organoid culture medium
Supplementary Table 2. Key resources

### Supplementary Figures

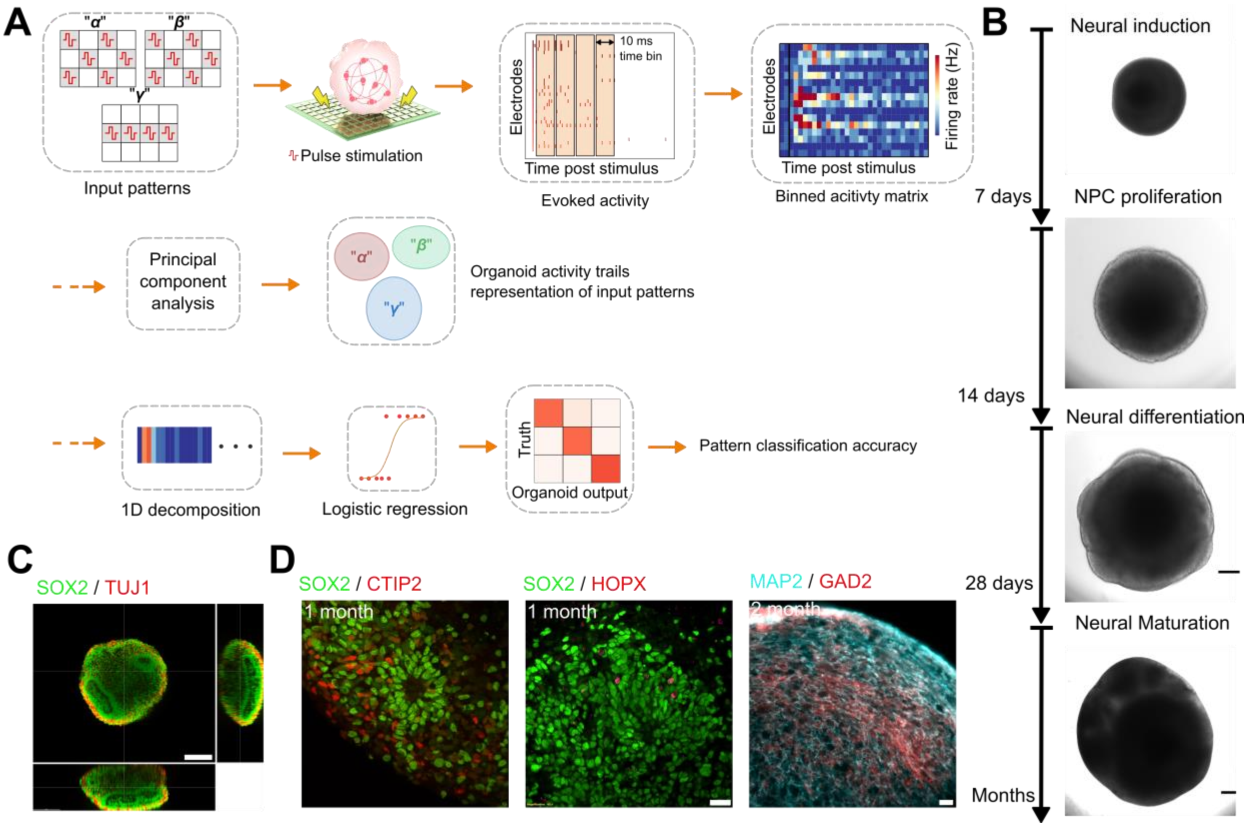

**Supplementary Fig.1. Workflow of functional phenotyping of brain organoids based on pattern classification. (Corresponding to Fig.1)**

(**A**) Three distinct input patterns were transformed into spatially patterned bipolar electrical stimulations (500 mV, 500 μm). Following stimulation, the evoked activity in brain organoids was recorded and converted into an activity matrix (MxN, where M represents the number of electrodes and N represents the number of 10-ms time bins). Each stimulation pattern was applied to the organoids 30 times, resulting in a total of 60 activity matrices for downstream principal component analysis (PCA) visualization and logistic regression, used to determine pattern classification accuracy.

(**B**) Overview of the cortical organoid culture protocol and representative organoid images of different time stages (Scale bar, 200 μm).

(**C**) Section images of whole-mount staining results, corresponding to **Fig.1C** (Scale bar, 100 μm).

(**D**) Representative immunofluorescence staining images of cortical organoids (Scale bar, 20 μm).

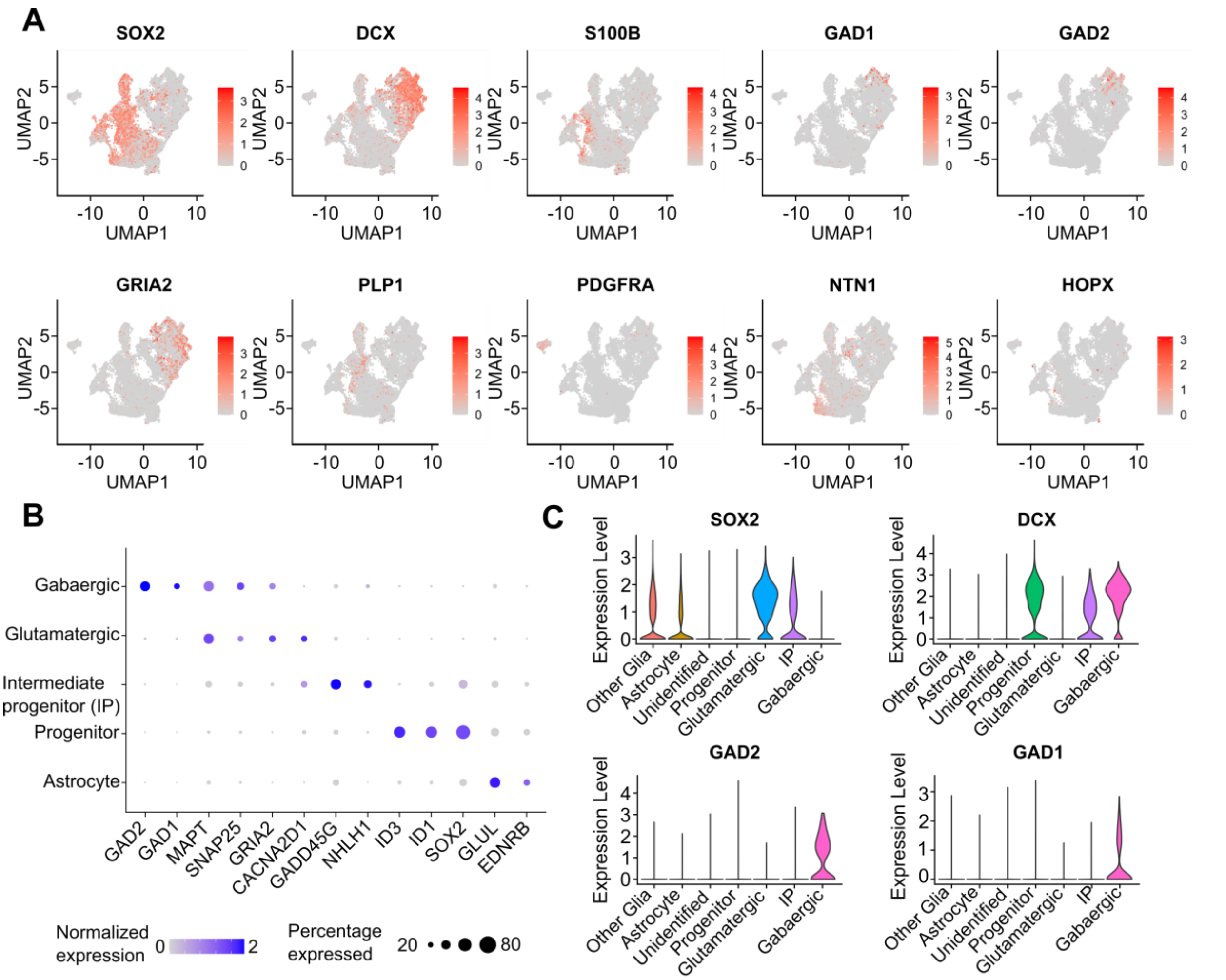

**Supplementary Fig.2. Single cell RNA seq data (Corresponding to Fig.1).**
(**A**) UMAP plots showing the expression levels of cell-type specific marker.
(**B**) Dot plots showing cluster-specific gene expression across the main cell clusters.
(**C**) Violin plots of progenitor marker (SOX2, DCX) and GABAergic neuron marker (GAD1, GAD2) gene expression across all clusters.

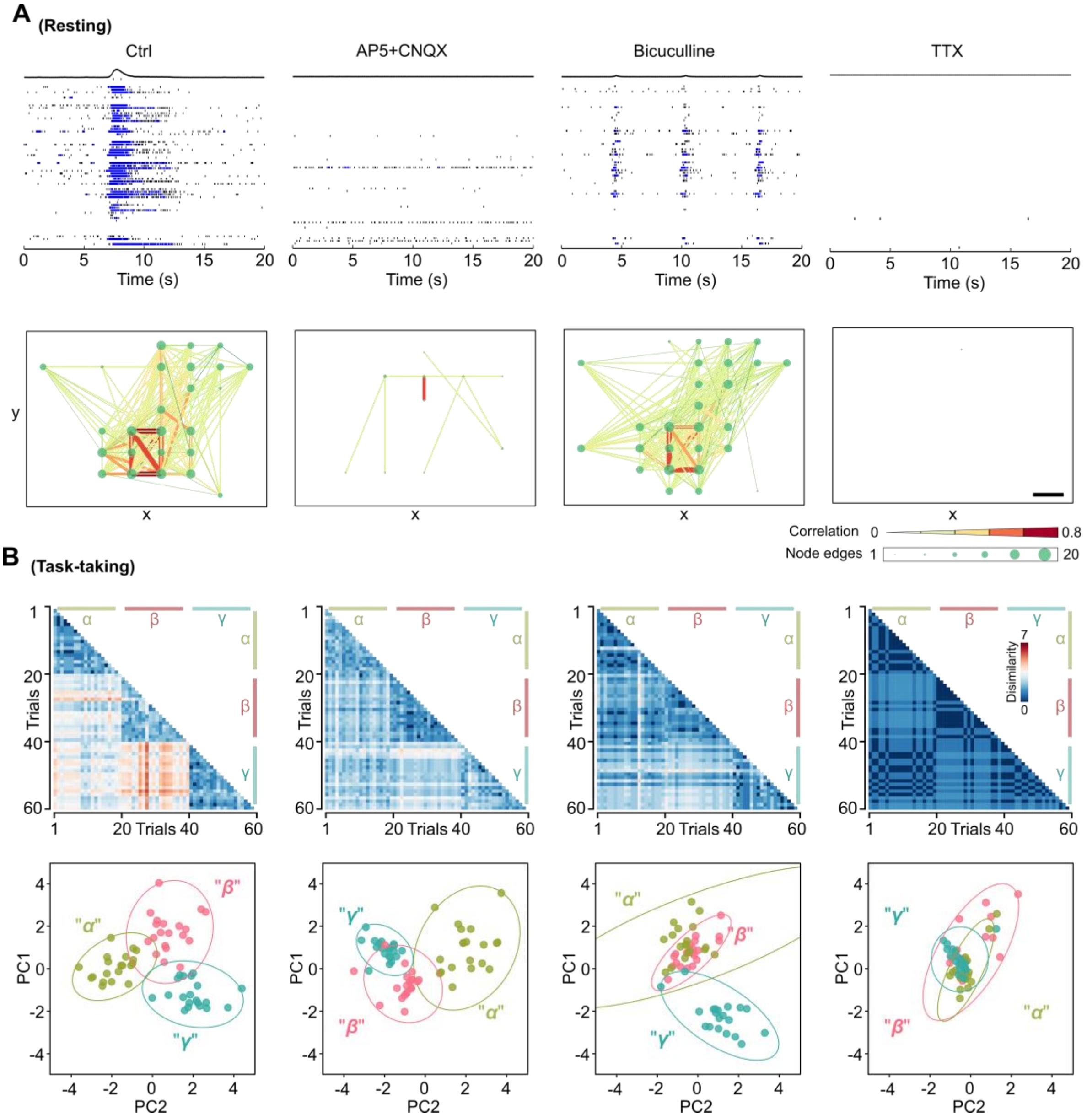

**Supplementary Fig.3. Characterization of organoid function changes by molecular neuromodulators. (Corresponding to Fig.3)**

(**A**) Resting-state MEA measurement of organoids following neuromodulator treatment. Top: Representative raster plots of organoid electrical activity in response to various neuromodulators—AP5+CNQX (AMPA and NMDA antagonists), bicuculline ($GABA_A$ antagonist), and TTX (tetrodotoxin). Bottom: Corresponding functional connectivity maps for organoids treated with each neuromodulator.

(**B**) Task-state MEA measurements of organoids treated with neuromodulators. Representative heatmaps show the dissimilarity (calculated by Euclidean distance) between each stimulation trial, along with principal component analysis (PCA) results depicting neural activity patterns in response to three distinct stimulation patterns across drug-treated organoids. The classification of these stimulation-induced activities reflects the pattern classification capability of the organoids.

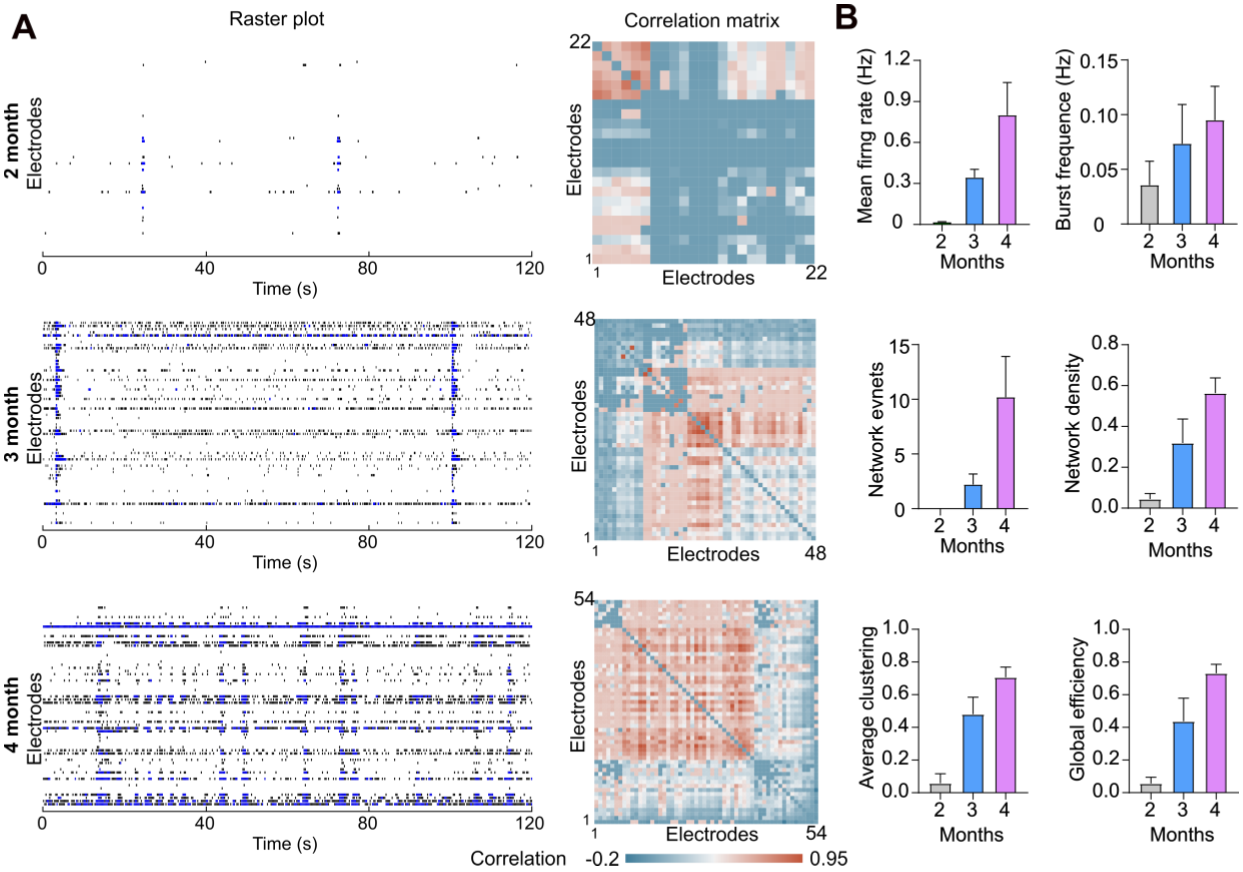

**Supplementary Fig.4. Maturation of functional neural networks in human cortical organoids. (Corresponding to Fig.3)**

(**A**) Functional maturation of cortical organoids. Representative raster plot and STTC correlation matrix of 2-, 3-, and 4-month-old human cortical organoids.

(**B**) Quantification results of spontaneous activities and functional connectivity in 2-, 3-, and 4-month-old human cortical organoids showing the functional maturation of human cortical organoids. (mean ± s.e.m., n = 5 organoids, 3 independent experiments).

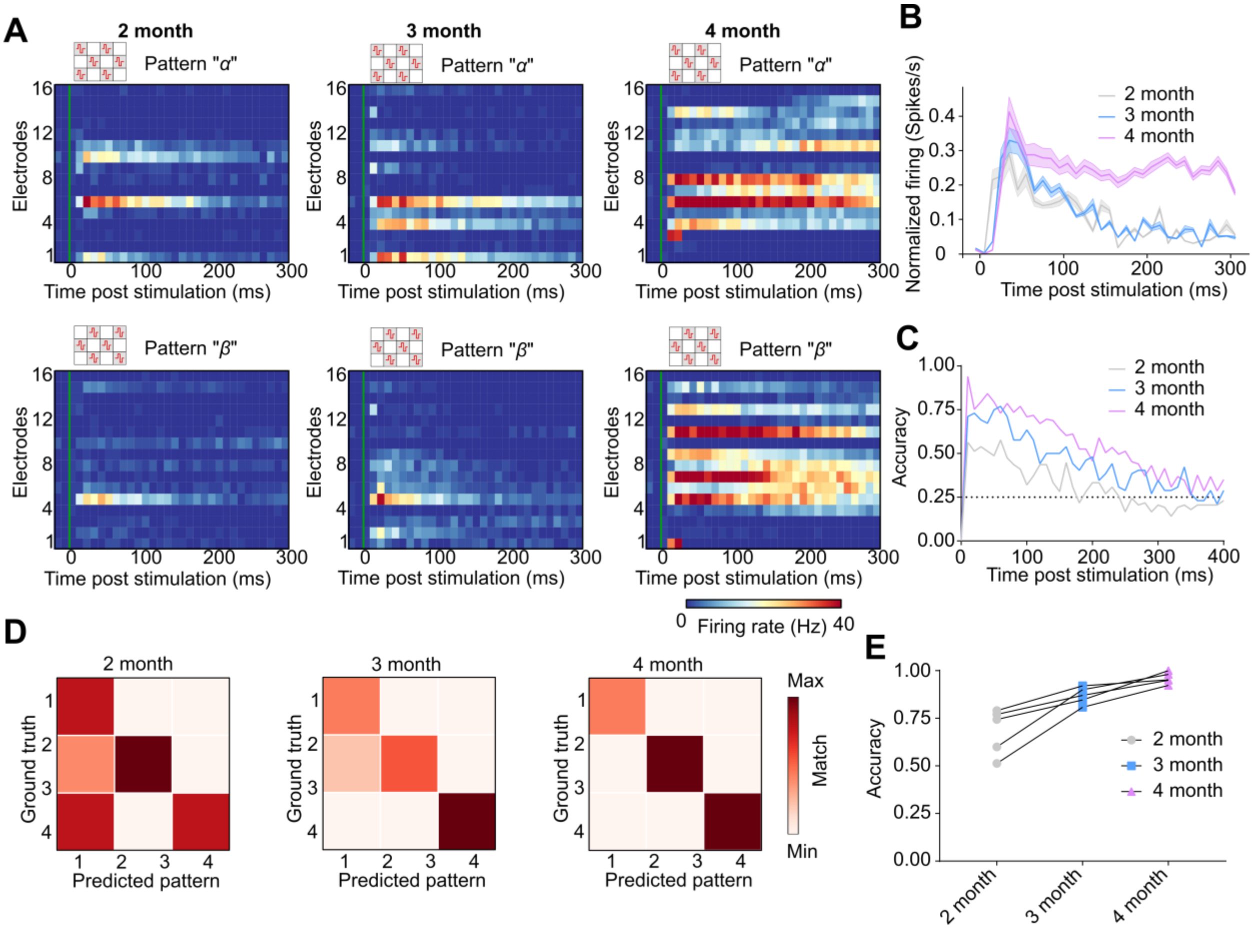

**Supplementary Fig.5. Functional phenotyping of developing cortical organoids. (Corresponding to Fig.3)**

(**A**) Representative raster plots of 2-month-old, 3-month-old, and 4-month-old organoids evoked by two complementary spatial patterns (Pattern 1 and 2) of electrical stimulation pulses.

(**B**) Post-stimulation histogram of 2-month, 3-month, and 4-month organoids evoked by pattern 1 stimulation (corresponding to **a**).

(**C**) Representative dynamics of 4 patterns classification accuracy post stimulation of 2-month-old, 3-month-old, and 4-month-old organoids.

(**D**) Representative confusion matrix heatmap of 3 patterns classification over 3-month maturation.

(**E**) 3 pattern recognition accuracy corresponding to cortical organoids development time (n=5, from 3 independent experiments).

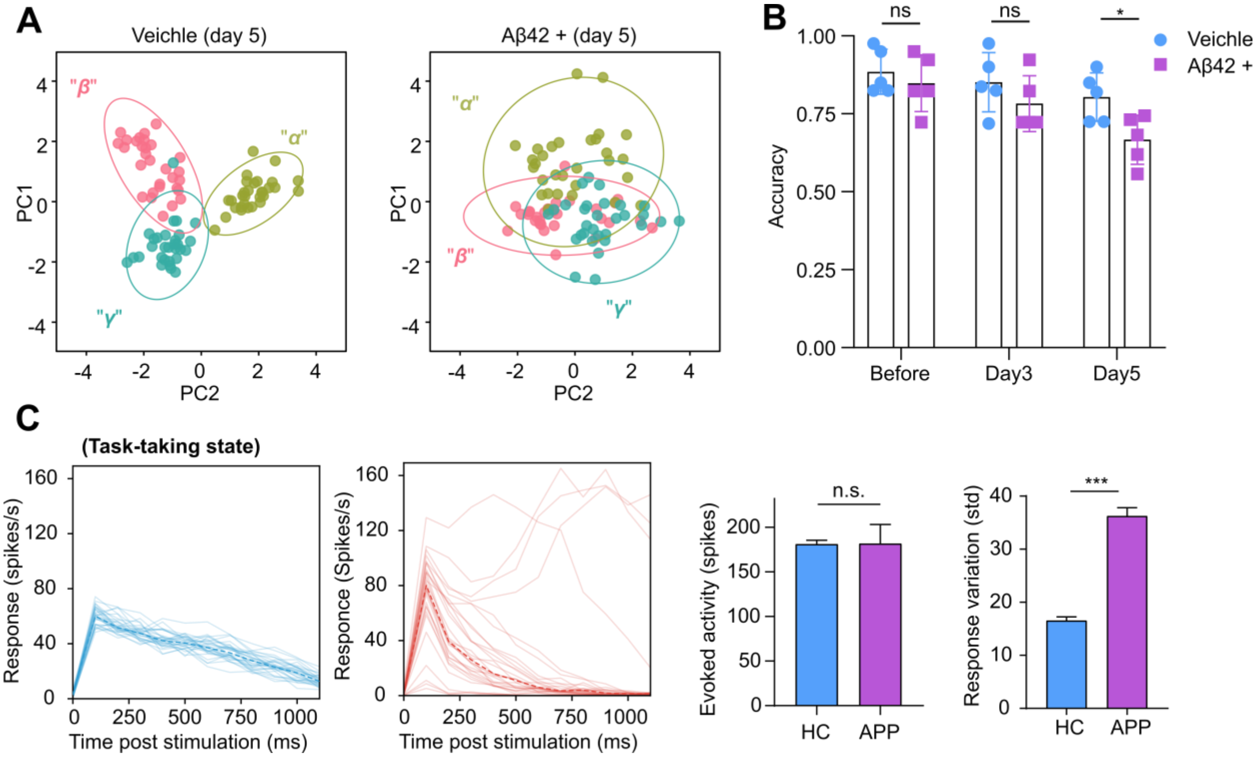

**Supplementary Fig.6. Functional phenotyping ONNs impacted by Aβ (Corresponding to Fig.4).**

(**A**) Representative principal component analysis (PCA) results of three patterned stimulation-induced neural activities in the Aβ42 + (treated with 1500pg/ml Aβ42) and vehicle organoids on day 5.

(**B**) Pattern classification accuracy of Aβ42 + and vehicle organoids over time of treatment. (mean ± s.e.m., n = 5, from 3 independent experiments).

(**C**) Overlayed peri-stimulation histogram of evoked activities in HC and APP cortical organoids stimulated with pattern $\alpha$, each line represents a trial, and the corresponding quantification of the standard deviation of evoked responses. corresponding to **Fig.4D.**

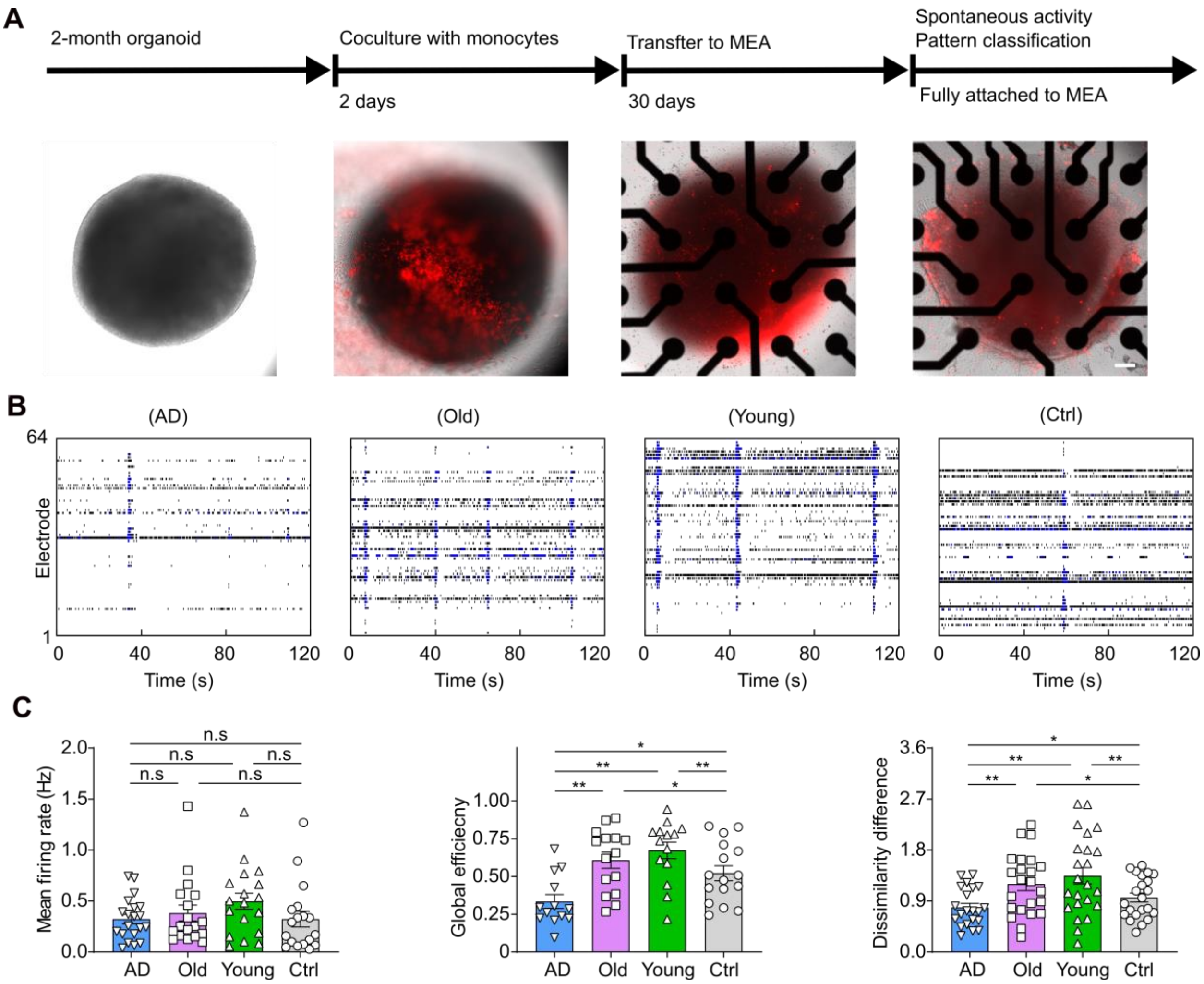

**Supplementary Fig.7. Monocytes impact organoid functional networks (Corresponding to Fig.5).**

(**A**) Workflow showing the functional phenotyping of organoids cocultured with monocytes. Scale bar = 100 μm.

(**B**) Representative raster plots showing the spontaneous activities of organoids under various conditions.

(**C**) Quantification of functional network (ONN) alterations in response to AD, old, and young monocytes, respectively (mean ± s.e.m., n = 15, 3 independent experiments).

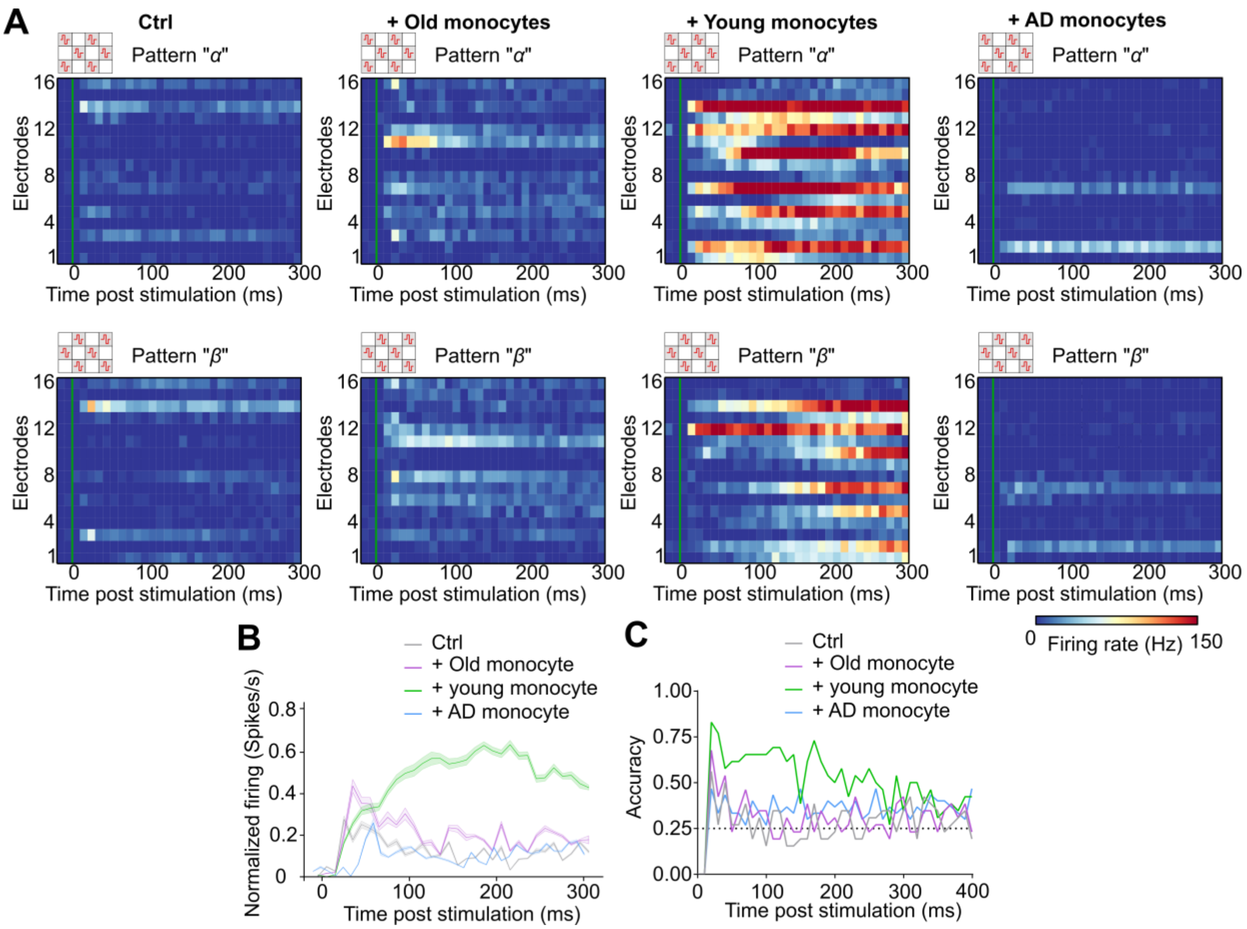

**Supplementary Fig.8. Organoid phenotype impacted by young, old, and AD monocytes. (Corresponding to Fig.5)**

(**A**) Representative raster plots of cortical organoids cocultured with young, old, and AD monocytes evoked by two complementary spatial patterns (Pattern $\alpha$ and Pattern $\beta$) of electrical stimulation pulses.

(**B**) Post-stimulation histogram of different group organoids responded to Pattern $\alpha$ (corresponding to **A**).

(**C**) Dynamic pattern classification accuracy in human cortical organoids treated by young, old, and AD monocytes, respectively (n=5, from 3 independent experiments).

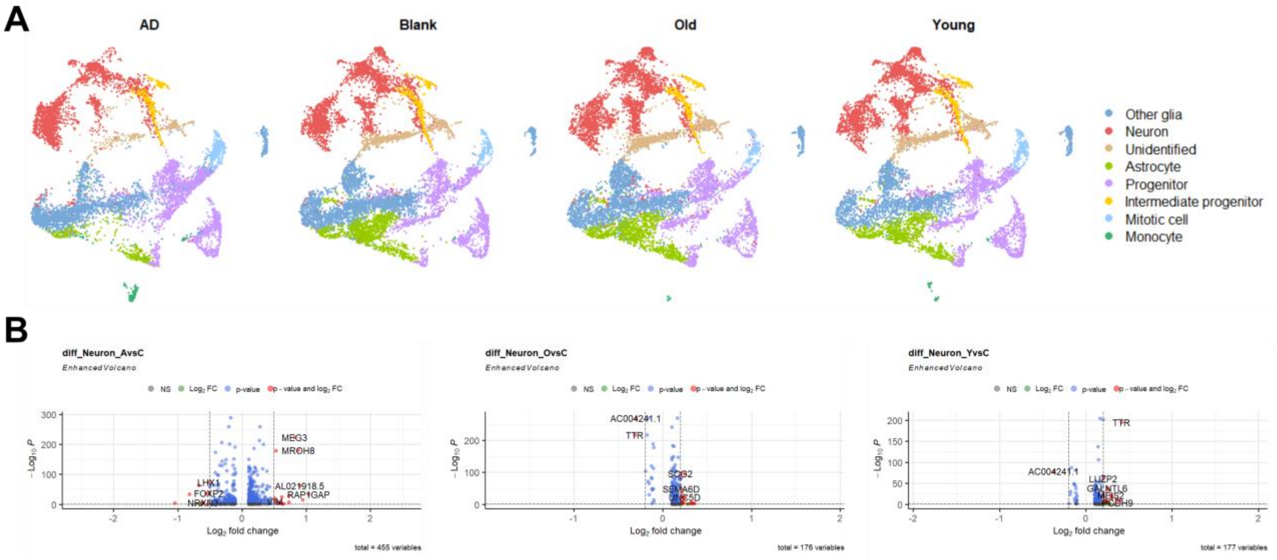

**Supplementary Fig.9. Additional single-cell seq data of monocyte-cocultured cortical organoids (Corresponding to Fig.5).**

(**A**) Uniform manifold approximation and projection (UMAP) plots of single-cell RNA-seq results from the integrated dataset (AD, Ctrl, Old, and Young), showing the clusters of various cell types within the organoids.

(**B**) Volcano plot showing the differently expressed genes in neurons among 'AD vs Ctrl', 'Old vs Ctrl', and 'Young vs Ctrl'.

**Supplementary Movie**

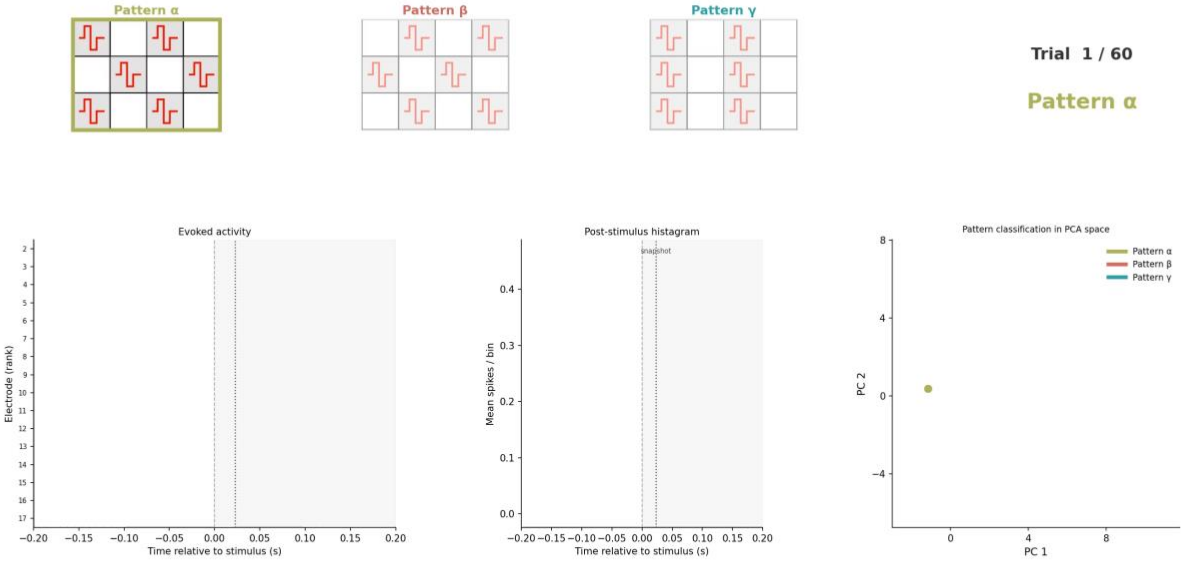

**Supplementary Movie 1. The pattern classification process on the Brainopheno platform.** Three electrical stimulation patterns (α, β, γ) were applied to a cortical organoid on the MEA (20 trials each). Top panel: The stimulation patterns and trial numbers. Bottom panel (Left to Right): Raster plot showing the evoked neural activity [-200ms, 200ms] of each trial. Post-stimulus histogram. Neural activity trajectory projected onto the PCA space; filled circles mark each trial's state at maximum separation (23 ms post-stimulus).

**Supplementary Table 1.**

| **Cortical Organoid Medium 1: EB Formation Medium** | **Day 0** |
|---|---|
| **COMPONENTS** | **CONCENTRATION** |
| mTeSR1 | 1X |
| SB431542 | 10 µM |
| XAV939 | 2 µM |
| Dorsomorphin | 1 µM |
| Y-27632 **(only for 24h)** | 90 µM |
| **Cortical Organoid Medium 2 - Proliferation Medium** | **Day 3** |
| **COMPONENTS** | **CONCENTRATION** |
| Neuralbasal | 1X |
| GlutaMax (100X) | 1X |
| Gem21 (50X) | 1X |
| N2 NeuroPlex (100X) | 1X |
| MEM-NEAA (100X) | 1X |
| Penn/Strep (100X) | 1X |
| SB-431542 | 10 µM |
| XAV939 | 2 µM |
| Dorsomorphin | 1 µM |
| **Cortical Organoid Medium 3- Expansion Medium** | **Day 10** |
| **COMPONENTS** | **CONCENTRATION** |
| Neuralbasal | 1X |
| GlutaMax (100X) | 1X |
| Gem21 (50X) | 1X |
| N2 NeuroPlex (100X) | 1X |
| MEM-NEAA (100X) | 1X |
| Penn/Strep (100X) | 1X |
| FGF-2 | 20 ng/mL |
| **Cortical Organoid Medium 4 - Differentiation Medium** | **Day 17** |
| **COMPONENTS** | **CONCENTRATION** |
| Neuralbasal | 1X |
| GlutaMax (100X) | 1X |
| Gem21 (50X) | 1X |
| N2 NeuroPlex (100X) | 1X |

| | |
|---|---|
| MEM-NEAA (100X) | 1X |
| Penn/Strep (100X) | 1X |
| FGF-2 | 20 ng/mL |
| EGF | 20 ng/mL |
| **Cortical Organoid Medium 5 - Maturation Medium** | **Day 24** |
| **COMPONENTS** | **CONCENTRATION** |
| Neuralbasal | 1X |
| GlutaMax (100X) | 1X |
| Gem21 (50X) | 1X |
| N2 NeuroPlex (100X) | 1X |
| MEM-NEAA (100X) | 1X |
| Penn/Strep (100X) | 1X |
| BDNF (100µg/mL) | 10 ng/mL |
| GDNF (10mg/mL) | 10 ng/mL |
| NT3 (100µg/mL) | 10 ng/mL |
| Ascorbic Acid | 200 µM |
| dibutyryl-cAMP | 1000 µM |
| **Cortical Organoid Medium 6 - Maintenance Medium** | **Day 31** |
| **COMPONENTS** | **CONCENTRATION** |
| Neuralbasal | 1X |
| GlutaMax (100X) | 1X |
| Gem21 (50X) | 1X |
| N2 NeuroPlex (100X) | 1X |
| MEM-NEAA (100X) | 1X |
| Penn/Strep (100X) | 1X |
| **Cortical Organoid Medium 7 - BrainPhys Medium** | **Day 45** |
| **COMPONENTS** | **CONCENTRATION** |
| Brainphys Imaging Optimized Medium | 1X |
| NeuroCult SM1 supplement | 1X |
| GlutaMax (100X) | 1X |
| cAMP (10mM) | 100 µM |
| Penn/Strep (100X) | 1X |

**Supplementary Table 2.**

| REAGENT or RESOURCE | SOURCE | IDENTIFIER |
|---|---|---|
| **Antibodies** | | |
| Rat CTIP2 | Abcam | Cat# AB18465, |
| Rabbit GAD2 | Cell Signaling | Cat# 5843, |
| Guinea pig vGlut1 | Millipore | Cat# AB5905, |
| Rabbit HOPX | Millipore | HPA030180, |
| Rabbit Tuj1 | Cell Signaling | Cat# D71G9 RRID:AB_10694505 |
| Goat SOX2 | R&D | Cat# AF2018-SP RRID:AB_355110 |
| Rabbit GFAP | Agilent Dako | Cat# Z033429-2 |
| Chicken MAP2 | Millipore | Cat# AB5543, RRID:AB_571049 |
| Alexa 488-conjugated anti-Rat IgG antibody | Life technologies | Cat# A-21208, RRID:AB_141709 |
| Alexa 488-conjugated anti-Goat IgG antibody | Jackson | Cat# 705-545-003, RRID:AB_2340428 |
| Alexa 594-conjugated anti-Rabbit IgG antibody | Life technologies | Cat# A-21207, RRID:AB_141637 |
| Alexa 594-conjugated anti-Mouse IgG antibody | Life technologies | Cat# A32744, RRID:AB_2762826 |
| Alexa 647-conjugated anti-Chicken IgG antibody | Jackson | Cat# 703-605-155, RRID:AB_2340379 |
| Alexa 647-conjugated anti-Rabbit IgG antibody | Life technologies | Cat# A32795, RRID:AB_2762835 |
| **Chemicals, peptides, and recombinant proteins** | | |
| Dorsomorphin | R&D Systems | 3093 |
| SB431542 | Stemcell | 04-0010-10 |
| Y-27632 dihydrochloride | Sigma-Aldrich | 688000 |
| Poly(ethyleneimine) solution | Sigma-Aldrich | 408700 |
| Laminin | Sigma-Aldrich | L4544 |
| Recombinant Human bFGF | Peprotech | 100-18B |
| Recombinant Human EGF | Peprotech | AF-100-15 |
| Recombinant Human NT3 | Peprotech | 450-03 |
| Recombinant Human BDNF | Peprotech | 450-02 |
| Recombinant Human GDNF | Peprotech | 450-10 |
| L-Ascorbic acid | Sigma-Aldrich | A4403 |
| N6,20 -O-Dibutyryladenosine 30 ,50 -cyclic monophosphate sodium salt | Sigma-Aldrich | D0627 |
| **Biological samples** | | |
| Healthy human monocytes | Stemcell | 200-0167 |
| AD patient monocytes | IU health | |
| **Deposited data** | | |
| Single-cell RNA-seq data | This paper | GEO: GSE310391 Review token: kzkbgqeavdcbziz Link: https://www.ncbi.nlm.nih.gov/geo/query/acc.cgi?acc=GSE310391 |

| | | |
|---|---|---|
| **Experimental models: Cell lines** | | |
| Human: H9 embryonic stem cells | WiCell Research Institute | WA09 |
| Human: APP duplication IPSCs | WiCell Research Institute | UCSD241i-APP2-3 |
| Human: Healthy donor IPSCs | New York Stem Cell Foundation | 051179 |
| **Software and algorithms** | | |
| Axis Navigator v3.5.1 | Axion biosystem | N/A |
| Cellranger v6.1.2 | 10X genomics | https://github,com/10XGenomics/cellranger |
| GraphPad Prism v8.0.64 | GraphPad Software | https://www.graphpad.com/features |
| ImageJ | NIH | https://imagej.net/Fiji/Downloads |
| Python v3.9.18 | Python | https://www.python.org/ |
| Seurat v4.3.0 | Github | https://github.com/satijalab/seurat |
| R v4.1.0 | CRAN | https://ourcodingclub.github.io/tutorials/intro-to-r/ |
| Elephant v1.0.0 | NeuralEnsemble | https://neuralensemble.org/elephant/ |
| cellSens Dimension v 2.3 | Olympus | N/A |
| NetworkX v2.5.1 | NetworkX | https://networkx.org/ |
| Matlab R2020b | MathWorks | https://www.mathworks.com/products/matlab.html |
| COMSOL Multiphysics v5.3a | COMSOL | https://www.comsol.com/ |
| Adobe | Adobe Photoshop CC | https://www.adobe.com/products/photoshop.html |
| Inkscape v1.3.2 | Inkscape | https://inkscape.org/ |
